# Large retrospective WGS study describes three major sequence types of *S. aureus* in India and reveals two novel multi-drug resistant sub-lineages of *S. aureus* Clonal Complex 22

**DOI:** 10.1101/2022.06.21.496943

**Authors:** Monica I. Abrudan, Varun Shamanna, Akshatha Prasanna, Anthony Underwood, Silvia Argimón, Geetha Nagaraj, Sabrina Di Gregorio, Vandana Govindan, Ashwini Vasanth, Sravani Dharmavaram, Mihir Kekre, David M. Aanensen, K. L. Ravikumar

## Abstract

**Background:** *S. aureus* is a major pathogen in India, causing nosocomial infections, but little is known about its molecular epidemiology and mechanisms of resistance in hospital settings. Here, we use WGS to characterize 508 *S. aureus* clinical isolates collected across India and analyze them in a global context.

**Methods:** Whole-genome sequencing was performed on 508 clinical isolates of *S. aureus* collected from 17 sentinel sites across India between 2014 and 2019 with the Illumina platform. AMR genotypes were predicted using Staphopia. Isolates carrying novel SCC*mec* cassettes were further characterized using long-read sequencing. A temporal analysis of clonal complex (CC) 22 global isolates from 14 different studies was performed using BactDating.

**Results:** Sequencing results confirmed 478 isolates as *S. aureus*. ST22, ST772 & ST239 were the major clones identified. An in-depth analysis of the 175 CC22 Indian isolates identifies two novel ST22 MRSA clones, PVL+ and one harboring the *tsst-1* gene. Temporal analysis showed that these two ST22 clusters shared a common ancestor in the 1980s and they became widespread after the year 2000 in India. Analyzing these in a global context, we found evidence of transmission of the two Indian clones to other parts of the world.

**Conclusion:** Our study describes a large retrospective *S. aureus* sampled from India. By comparing the Indian isolates globally we show the evidence of the international transmission of ST22 Indian isolates. Even though the two of the major dominant clones (ST772 and ST239) using WGS have been reported, this is the first study that describes the third dominant clone (ST22) in India.

**Impact statement:** *Staphylococcus aureus* is an opportunistic pathogen listed as a high-priority pathogen by WHO. It is a leading cause of nosocomial infections in India and worldwide. Our study is the first study to describe the epidemiology of *S. aureus* in India with a large sample set of 478. Here we describe a collection of 478 *S. aureus* genomes, isolated from 17 sentinel sites in India, between 2014 and 2019. With the focus on understanding sequence types, AMR profiles, SCC*mec* types, and *spa* types and discuss these in the context of previous molecular studies on *S. aureus* conducted in India. We also conducted an in-depth analysis of the Clonal Complex 22 Indian isolates and we identified two novel ST22 MRSA clones, both PVL+ and one harboring the *tsst-1* gene. Temporal analysis shows that these two ST22 clusters originated around 2010 in India. Analyzing these in a global context, we found evidence of transmission of the two Indian clones in other parts of the world. Analysis of a cluster of 33 isolates belonging to ST239 from a single hospital in Bangalore indicates an outbreak that persisted over the period of three years from a single contamination source. The novel SCC*mec* types identified in our study are characterized using long reads to understand their genetic structure.

**Data Summary:**

1. Illumina read files of the strains used in the study have been deposited in European Nucleotide Archive, BioProject PRJEB29740 (https://www.ebi.ac.uk/ena/browser/view/PRJEB29740?show=reads). A full list of accession numbers for all sequence read files is provided in Supplementary table 2.
2. Nanopore reads are submitted to ENA under the BioProject PRJEB50484.
3. Metadata and other related information on the strains are provided in the microreact project with different views in this link microreact.org/s.aureus_ghru_analysis.
4. Strain information for the ST22 samples used from other studies is provided in microreact at this link: https://microreact.org/project/2xDvKQhriNveJ4kiVYsmSQ-s-aureus-wgs-study. The authors confirm all supporting data, code and protocols have been provided within the article or the supporting data repository.

## Introduction

*Staphylococcus aureus* is an opportunistic pathogen and a leading cause of community and nosocomial infections, which has prompted WHO to include it on a list of priority pathogens list for R&D of new antibiotics in 2018 (1). Methicillin-resistant *S. aureus* (MRSA) originally appeared in hospitals in the 1960s and then re-emerged in the community and hospitals in the 1980s, spreading worldwide and creating reservoirs in both settings. Traditionally, hospital-acquired MRSA (HA-MRSA) and community-acquired MRSA (CA-MRSA) were considered different clones, as they used to differ in their antibiotic susceptibility pattern and their mode of acquisition. Moreover, while HA-MRSA mostly affected certain at-risk populations in hospitals, CA-MRSA caused serious illness or even death to otherwise healthy individuals (2). The situation has changed in recent years when community-acquired MRSA (CA-MRSA) has become more invasive and transmissible and increasingly difficult to differentiate from HA-MRSA.

### *S. aureus* in India

Reports from the second half of the 1990s show that MRSA was endemic in Indian hospitals, and in some places, more than 50% of the hospital staff were found to harbor MRSA at any given point (3). The first genotypic study of *S. aureus* from India, which was conducted between 2003 and 2004 in Bangalore (4), and a typing study of random isolates from the Asian Bacterial Bank collected between 1998-2003, both identified international clones ST239 and ST241 as the major circulating sequence types in hospitals at the time. Subsequent larger studies, conducted after 2004 across multiple sites, found a higher *S. aureus* sequence type diversity, which included ST22, ST772, ST291, as well as ST239 (5) (6) (7) (8).

Recent evidence confirmed that ST22, ST772, and ST239 were the major MRSA STs in India (9) (10) (11), but other STs were also gaining importance, such as CA-MRSA ST2371, which was first identified in Indian hospitals in a tertiary care hospital in Mysore, South India, in a study from 2012-2013 (12). Out of these, ST22-MRSA-IV, ST772-MRSA-V, ST672-MRSA-V (9), ST8-MRSA-IV, and ST2371-MRSA-IV (a single locus variant of ST22) (12) have often been mentioned as being CA-MRSA and described as carrying a smaller *Staphylococcal* cassette chromosome *mec* (*SSCmec*) type (IV, V, or VII) (13) and the Panton-Valentine leukocidin (PVL toxins), a bicomponent leukocidin encoded by the *lukS-PV* and *lukF-PV* genes, residing in a prophage. PVL toxin causes leukocyte destruction, tissue necrosis, and increased disease severity and it is thought that they contribute to the success of some CA-MRSA lineages (13) (14).

### Dominant sequence types in India

#### ST772, or the “Bengal Bay” clone

ST772 is now one of the dominant sequence types in India and it is thought to have emerged in the Indian subcontinent during the early 1960s (15). It was suggested that the acquisition of SCC*mec* V (5C2) and the double serine mutations (S84L, S80Y) in genes *gyrA* & *grlA* leading to ciprofloxacin resistance have resulted in the expansion strains leading to the successful survival of multi-drug resistant ST772 in hospital settings in India (16) (17), but also in other parts of the world (15). ST772-MRSA-V clones are characterized by the presence of a prophage ΦIND772 with PVL and the staphylococcal enterotoxin A (*sea*), three distinct pathogenicity islands (vSa-alpha, beta, and gamma), and an integrated resistance plasmid (IRP), encoding a cluster of resistant genes such as *blaZ* (conferring resistance to Beta-lactams), *mphC* and *msrA* (conferring resistance to Macrolides), *aphA III* and a partial *aadE* (conferring resistance to Aminoglycosides) and *sat4* (conferring resistance to Streptothricin) (11).

#### In India, ST239 is slowly being replaced by ST22 and ST722

The ST239-MRSA is an invasive, highly recombinant, and virulent clone that causes a wide range of life-threatening infections. It has caused hospital epidemics starting from the 1970s throughout the world (18). It is still one of the dominant clones in Asia (19) and in Southern India (20) (21), although it is slowly being replaced by ST22 and ST722 (7). ST239-MRSA clones were found to carry several types of the SCC*mec* cassette, including SCC*mec* type V, III, IV, and I (22). A multi-drug resistant ST239-MRSA clone has been reported in India in several studies since 2010, which is known to carry the SCC*mec* cassette type III and the PVL genes, and are highly resistant to mupirocin and also has inducible clindamycin resistance (22).

#### Knowledge gap about the Indian Clonal Complex 22

*S. aureus* ST22-MRSA gained international attention in the late 80s when the ST22 sub-clone EMRSA-15 began spreading in English hospitals and soon after that, all over the world. Some of the important characteristics of EMRSA-15 were resistance to Fluoroquinolone and Macrolides and its small SCC*mec* cassette (type IVh) (23). More recently, another successful ST22-MRSA lineage, genetically distant to the EMRSA-15, was identified in communities of healthy children and adults in the Gaza Strip (24), and later in Russia (25). It was characterized by the SCC*mec* cassette type IVa, and an arsenal of toxin genes, including *chp* (CHIPS), *scn* (SCIN), *sak* (staphylokinase), and the *tsst-1* (toxic shock syndrome toxin-1). In the past years, the third lineage of ST22-MRSA, CA, and PVL+, has been reported in Asia and the Middle East, in countries such as India (26), Kuwait (27), Nepal (28), Japan (29), China (30) and Saudi Arabia (31). Although many studies initially assigned the CA-MRSA PVL+ to the European EMRSA-15 ST22, further analysis of SCC*mec* subtypes revealed that this clone originated most likely in Asia and was different from the European one (31). To our knowledge, the dominant *S. aureus* ST22-MRSA identified in Indian hospitals by (26), (7), (5) (6) has not yet been analyzed using the whole-genome sequencing, and the current paper aims to give an overview of the history, genomics, and epidemiology of this very important clone, which has been previously wrongly assigned to EMRSA-15.

#### The aims of this study

Here we characterize the Indian *S. aureus* population based on a large WGS retrospective surveillance study of 478 clinical isolates, collected from 17 sentinel sites across 10 states in India, between 2014 and 2019. Moreover, we contextualized the 175 clonal complex (CC) 22 isolates (out of which, 147 belong to ST22) from India with a set of 1624 CC22 isolates, from publicly available collections. Although typing studies are helpful for understanding the current make-up of dominant clones, whole-genome sequencing (WGS) provides a richer resource for surveillance studies and identification of local outbreaks and facilitates a comprehensive understanding of specific microbes’ dynamics.

## Results

### Population structure of S. aureus across 17 sentinel sites in India

To characterize the population structure of *S. aureus* in India, 17 sentinel sites were asked to contribute with MRSA isolates collected from patients between 2014 and 2019, to be sent for whole-genome sequencing.

509 of the received samples were confirmed as *S. aureus* with Vitek-2 (AST card P628, Biomerieux). The WGS of these isolates confirmed (n=478) as *S. aureus*, using *in silico* methods. 85 isolates out of the 478 were later confirmed as MSSAs, using phenotypic and genotypic methods, and they were also included in the analysis. The samples collected were from patients varying from the age range of 4 days to 100 years, comprising 65.89 % (n=315) male and 34.1 % (n=163) female. The specimen sources were mostly from pus (n= 274), followed by wound (n=43), tracheal (n= 37) and blood (n=19). Details about data distribution per year can be found in *Supplementary table 1*.

WGS showed that the strains belonged to 37 different sequence types (STs) and 3 other novel STs. ST22 (n=138), ST239 (n=72) and ST772 (n=66) were found to be the predominant STs. MRSA isolates mostly belonged to ST22 (n= 135), ST239 (n=72) and ST772 (n= 59), whereas MSSA isolates belonged to ST1 (n= 11), ST672 (n=9), ST2066 (n= 9), ST291(n= 7), and ST772 (n=7). The varied diversity of 90 different *spa* types was identified through *spa* typing. The dominant *spa* type was *spa-t852*, which was present solely in CC22-MRSA isolates (n=83).

A phylogeographic analysis of the whole collection was performed as described in the Methods section and this revealed that the major STs (ST22, ST239 and ST772) were present across the entire country.

### Antimicrobial Resistance

We performed susceptibility tests of the isolates for 15 antibiotics representing 11 antibiotic classes using Vitek-2 compact system (Biomerieux). All isolates showed susceptibility to Vancomycin and Teicoplanin, and most isolates were resistant to Penicillin (n=470), Ciprofloxacin (n=463), and Oxacillin (n=393). WGS AMR profiling was performed on 10 antibiotic classes and the resistance determinants are shown in more detail in Supplementary Table 2.

### ST772 is a major sequence type in our collection

13.8% of the total isolates (66/478) from our study were ST772, and they were spread across 10 sentinel sites throughout India. 59 of them were MRSA, out of which 86% (n = 51) carried an inverted version of the SCC*mec* V (5C2), and contained an extra *aac*(6’) gene, flanked by the insert sequence IS256 (Supplementary Figure 3). Another seven MRSA isolates carried the composite SCC*mec* V (5C2 and 5) cassette and one isolate was found to carry SCC*mec* IVc. We identified Φ IND772 prophage carrying the PVL operon (*lukS* and *lukF*) and a staphylococcal enterotoxin A (*sea*) in 53 of the isolates (see Methods). Co-location of *lukS, lukF* and *sea* genes on the Φ IND772 prophage was confirmed using Nanopore sequencing (see Methods).

### Identification of a potential persistent ST239 outbreak in one Indian sentinel site, using WGS

72 isolates in the study (15%) were found to be ST239. They were isolated from five sentinel sites across the country and 100% (72) of the isolates were MRSA. *In silico* analysis predicts resistance to Beta lactams (72), Macrolides (52), Aminoglycosides(70), Quinolones (72,), Mupirocin(3), Fusidic Acid(72), Fosfomycin(72), Tetracycline (72), Trimethoprim (19) and Rifampicin (56). All of the ST239 isolates carry a composite version of the SCC*mec* cassette type III. We further investigated the representative sample of the ST239 (G18252308, see Supplementary Figure 2) by Nanopore sequencing. This isolate has an 83% SCC*mec* template coverage (template sequence AB037671.1), according to SCC*mec*Finder 1.2. The mer operon, responsible for resistance to mercury & pT181has been lost in these samples probably mediated by IS431 transposition/recombination. Also, this SCC*mec* from India carries extra genes previously found in SCC*mec* type I. Coppens previously described this element in 2018 (32), but the current SCC*mec* found in G18252308 has a variant of the pls gene.

We have observed a cluster of 33 isolates from a single hospital in Bangalore, collected between 2014 and 2016, as shown in Figure 1B. A detailed investigation (see Methods section) showed that the mean pairwise SNP difference between the 33 isolates was 7 (range 0-18), indicating an outbreak that persisted over the period of three years. The mean pairwise SNP difference between isolates outside of this cluster was 82 (range 0-173). The star-like phylogeny of the 33 isolates indicates the existence of a single contamination source and the absence of person-to-person transmission. One study from India showed that 50% of staff harbored MRSA at any given point (33), which makes it plausible to believe that the single contamination source could be a healthcare professional, as in the case of the MRSA outbreak in Cambridge (34).

**Figure 1.**
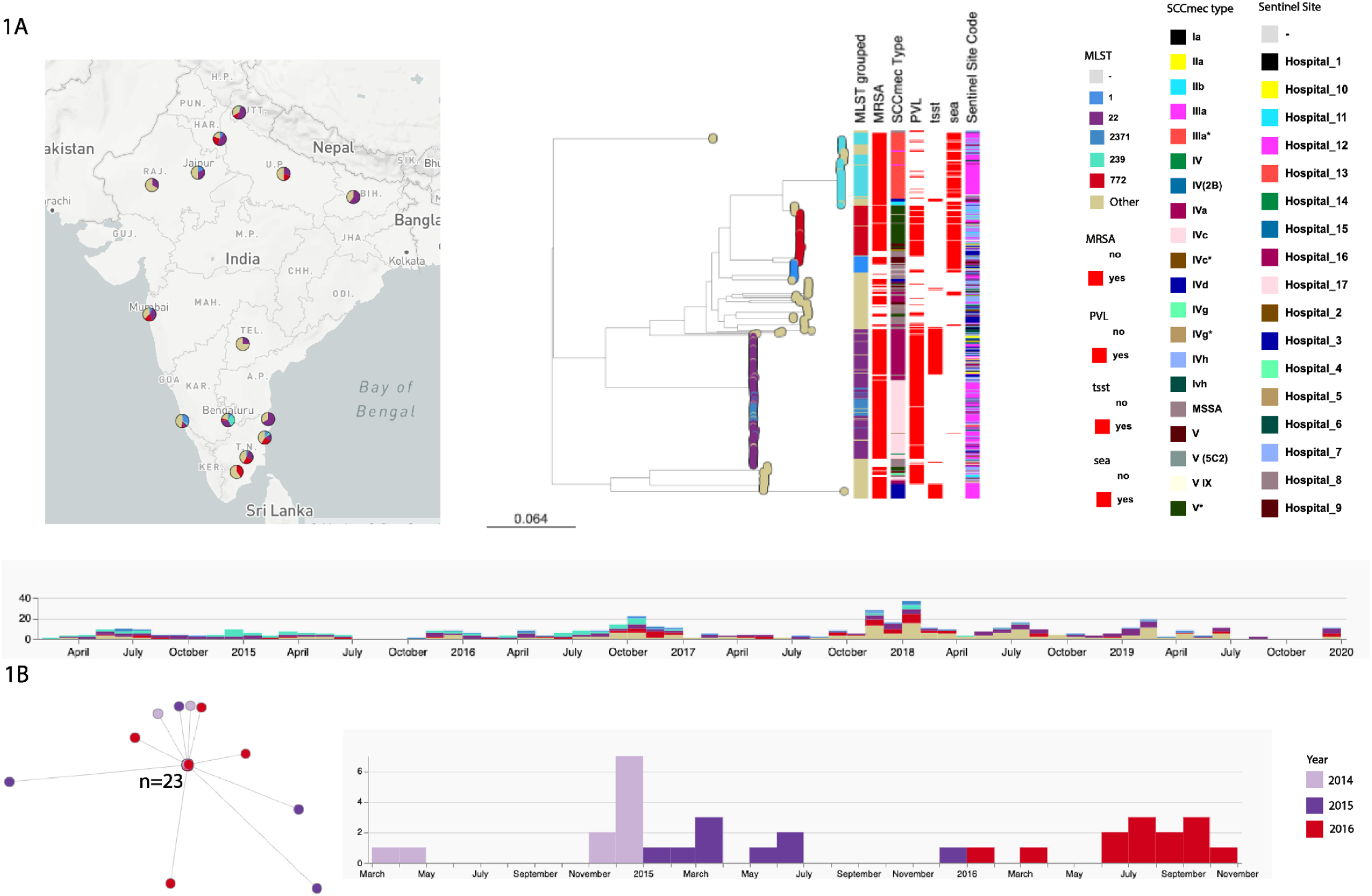
A. Phylogeny, molecular and demographic characteristics of the 478 Indian S. aureus isolates. The midpoint-rooted phylogenetic tree was inferred from 97,120 informative sites obtained after mapping the genomes to the complete genome of strain *Staphylococcus aureus* MSSA476 (strain GCF_000011525.1 (ST1)) and masking regions of recombination and MGEs. Tree nodes are colored according to their sequence types and are indicated on the map. Scale bars represent the number of single nucleotide polymorphisms (SNPs) per variable site. This view is available at: https://microreact.org/project/2xDvKQhriNveJ4kiVYsmSQ-s-aureus-wgs-study#i6az-ghru-tree B. A detailed view of the 33 ST239 outbreak isolates collected between 2014 and 2016 from Bangalore. This view is available at: https://microreact.org/project/2xDvKQhriNveJ4kiVYsmSQ-s-aureus-wgs-study#yxiv-st239-outbreak

**Figure 2.**
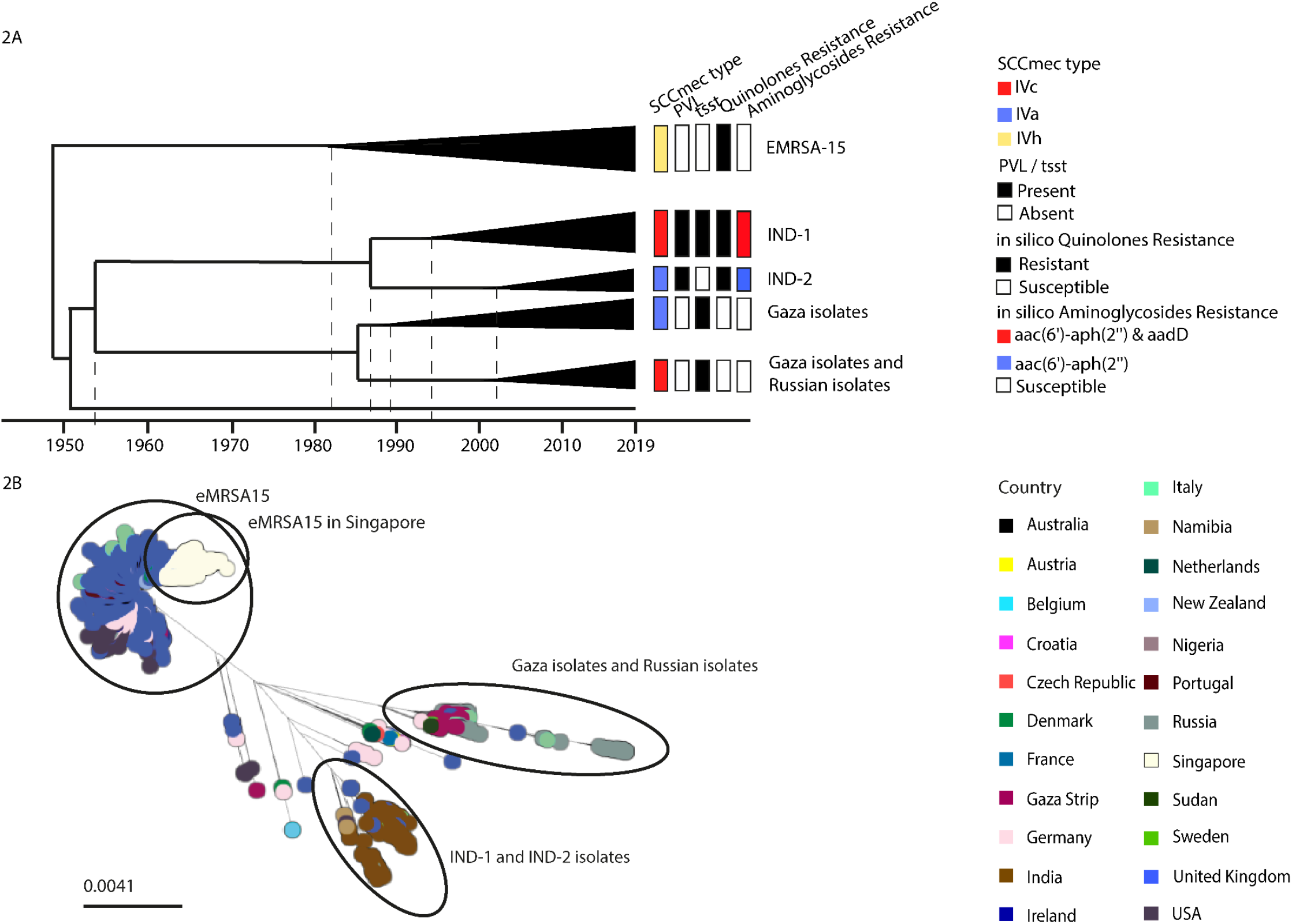
Indian CC22 strains in a global context. (A) A simplified temporal tree of the ST22 global collection; The data used to generate this tree is available at https://microreact.org/project/globalCC22. The metadata columns show the SCCmec types (based on the Staphopia predictions), presence/absence of the PVL and tsst genes and the in silico predicted resistances to Quinolones and Aminoglycosides. (B) Phylogenetic tree of the ST22 global collection. The midpoint-rooted phylogenetic tree was obtained from 44,703 SNPs, after mapping the genomes to the complete genome of strain EMRSA_15_GCF_000695215 and masking regions of recombination and MGEs. Tree tips are colored by country of origin as described in the legend. https://microreact.org/project/2xDvKQhriNveJ4kiVYsmSQ-s-aureus-wgs-study#6w7r-st22-gobal-view Scale bars represent the number of single nucleotide polymorphisms (SNPs) per variable site.

### Two novel ST22 MRSA clones, both PVL+ and one harboring the tsst-1 gene

175 (36%) isolates from the current GHRU collection belong to clonal complex 22 (CC22), out of which, 138 belong to sequence type 22 (ST22), 26 to ST2371, and 11 to other STs. The CC22 isolates were predicted, using in silico methods, to be resistant to Beta lactams (138), Macrolides (84), Aminoglycosides (135), Quinolones (137), Tetracycline (5), Trimethoprim (122). 172 isolates carried SCC*mec* IV and the predominant subtypes were IVc (n=105), and IVa (n=66). Three isolates were MSSA. The Indian CC22 isolates form two distinct clusters, supported by the bootstrap values.

### A temporal analysis of the CC22, using a large collection of global isolates, shows that the CC22 Indian isolates are from one monophyletic group, consisting of two distinct sub-clusters (IND-1 and IND-2), with an MRCA around the 2000s

In order to better understand the characteristics of the Indian isolates, we created a large collection of 1798 ST22 isolates from previously published studies (1623 ST22 global isolates & 175 CC22 isolates from this study), out of which, 1408 belonged to the EMRSA-15 clone and 105 to the ‘Gaza clone’. A temporal analysis using BactDating (35) indicates that the IND-1 clone became widespread around the year 2000, similar to the IND-2 clone, and the two share a common ancestor in the 1980s. Moreover, ST2371 emerged as an SLV of IND-1 around the year 2005. The analysis of the global collection also reveals that clade IND-2 includes an isolate from St. Petersburg, Russia, published recently by (36) (isolate SRR12560298, referred to in the paper as RU 1246, 2019, St. Petersburg), showing evidence of international transmission. The isolate from St. Petersburg was reported as having the highest number of pairwise SNPs among the other isolates in the Russian collection, and carried virulence genes: *sec, sell, tsst-1, fnbA, fnbB*, and also the SCC*mec* IVa cassettes which were also seen in all of the Indian isolates of this study.

### The CC22 IND-1 Cluster

The first cluster, which we called IND-1, is formed of 107 PVL+ isolates, identified in 9 of the 17 contributing sentinel sites. The average pairwise SNP difference between IND-1 isolates is 86 (0-195) and based on *in silico* methods, these isolates were predicted to be resistant to Beta lactams (107), Macrolides (52), Aminoglycosides(106), Quinolones (106), Tetracycline (2) and Trimethoprim (86).

An analysis of the accessory genomes of Indian ST22 isolates using Panaroo (37) revealed that all genomes in the IND-1 cluster carry the sraP gene. We found that the genomes in the IND-1 cluster carry a 111 amino acid shorter version of the SraP protein, compared to the SraP version of the protein found in the isolates forming the IND-2 cluster. SraP is a Serine-rich adhesin for platelets, which mediates binding to human platelets, possibly through a receptor-ligand interaction, and is a potential virulence determinant in endovascular infection (38). Clade-specific versions of the SraP proteins were also identified in a study of the CC30-MRSA from Argentina (39).

The SCC*mec* cassette type IVc found in the IND-1 isolates is 80% similar to the SCC*mec* cassette AB096217 (in Genbank). Using long-read sequencing (see Methods), we established that the current version of the SCC*mec* type IVc had an integration of puB110 plasmid which carries the *aadD1* & *bleo* aminoglycoside resistance genes (see Supplementary Figure) and that the *aacA-aph* gene, which shows on the AB096217 reference sequence, has translocated into a different location in the IND-1 genomes.

### ST2371, a sub-clone of the CC22 IND-1 cluster

ST2371 is a single locus variant (SVL) of ST22 and is a prevalent community-associated MRSA, multidrug-resistant clone, which was previously reported in Southern India (12) (8), but also sporadically in other parts of the world (40). The 26 ST2371 Indian isolates identified in this study were collected from four sentinel sites and are closely related to the other ST22 isolates from the IND-I clade (mean pairwise SNP difference between the ST22 and the ST2371 sub-clades is 85 (min 69, max 173)). Similar to the 58 ST2371 isolates from a 2011 outbreak in a neonatal unit in Cambridge UK (34), the ST2371 Indian isolates identified in this study carry the SCC*mec* cassette type IVc and are PVL+. Also, the mean SNP difference between the Cambridge ST2371 isolates and the Indian ST2371 isolates is 54 (min 36, max 76)), which suggests that the Cambridge lineage likely originated from India.

### The CC22 IND-2 Cluster

The second Indian cluster, which we called IND-2, comprises 61 isolates collected from 10 sentinel sites, with an average pairwise SNP difference between isolates of 96 (0 - 359). Unlike isolates from clade IND-1, isolates from clade IND-2 carry the SCC*mec* cassette type IVa, are susceptible to Aminoglycosides, and harbor the *tsst1* gene. An analysis of the ST22 pangenome, identified the genes *entC1, gloB* (*uncharacterized Metallo-**β**-lactams*), and *tsst* as being specific to the genomes in the IND-2 cluster (see Methods section). Enterotoxin C, a product of *entC1*, is a heat-stable enterotoxin produced by *S. aureus*, that activates the host immune system by binding as unprocessed molecules to major histocompatibility (MHC) complex class II and T-cell receptor (TCR) molecules. *entC1* is often found in processed food from raw milk, can cause food poisoning and its presence in dairy products has been used as an indicator of food contamination (41).

Enterotoxin C (42) although absent from the eMRSA reference genome, was also present in the majority of the ST22 eMRSA genomes (the clade originally called ST22-A), in a study by (23). When compared to the reference genome EMRSA-15, the ST22 isolates from the IND-2 cluster have a shorter version of the IS200-like transposase at position 141546 (a decrease of 255 nucleotides, from 486 to 231). Also, the same isolates have shorter *ComF* operon protein 1 than the reference at position 789971, by 273 nucleotides. In silico analysis predicts resistance to Beta lactams (61), Macrolides (49), Aminoglycosides(61), Quinolones (61), Mupirocin(1), Tetracycline (1), and Trimethoprim (61).

The pangenome analysis using panaroo (37) performed for global ST22 isolates without the isolates from this study showed 2192 cores genes and 1292 accessory genes. When the CC2 isolates from this study were added to the global set the core genes increased to 2200 & accessory genes to 1406. The detailed results of the pangenome results are given in supplementary table 2.

## Discussion

The present study offers an unprecedented view of the MRSA in India from isolates collected between 2014 and 2019 from 17 sentinel sites. Earlier studies have described two of the major dominant clones (ST772 and ST239) using WGS, and this is the first study that describes the third dominant clone (ST22) using the WGS approach. In addition, the current study also offers details of the less studied sequence types and provides a clear understanding of phylogenetic relationships between these. 15 new sequence types were found and submitted to PubMLST for identification.

An in-depth analysis of the ST22 Indian isolates identified two novel ST22 MRSA sub-lineages, both PVL positive and one harboring the tsst-1 gene. Temporal analysis showed that these two ST22 clusters originated around 2000 in India. An analysis of these in a global context found evidence of transmission of the two Indian clones to other parts of the world. The study shows evidence of the international transmission of isolates in the IND-1 cluster to the UK and to Italy and of isolates in the IND-2 cluster to Russia (see the Microreact view microreact.org/project/ghru-st22-isolates-in-global-context). In the absence of WGS data, several previous studies have misidentified ST22 isolates as eMRSA15, however, we were able to show that isolates with similar resistance and virulence profiles belong to the newly identified Indian lineages. The study also gives a comprehensive view of the ST2371 as a new emerging lineage in India, a sublineage of CC22, and describes it in relationship with the other ST22 isolates.

We have observed a cluster of 33 isolates from a single hospital in Bangalore, and a detailed investigation indicated an outbreak that persisted over the period of three years from a single contamination source. Implementation of routine surveillance and simple intervention measures are the need of the hour to reduce these outbreaks (43).

For selected isolates, we were able to produce single-contig assemblies through Nanopore sequencing to present the structure of three SCC*mec* cassette types. This was not achieved using short-read sequencing and known typing software tools, such as SCC*mec*Finder and Staphopia. This enabled us to describe the SCC*mec* types not previously reported such as the SCC*mec* cassette found in ST772 isolate belonging to type V carrying the aac(6’) gene, which confers resistance to Aminoglycosides. ST22, IND-1 cluster, belonging to SCC*mec* type IVc carries an extra insert sequence with genes aadD1 and bleO conferring resistance to Aminoglycosides and Bleomycin respectively. SCC*mec* cassette found in ST239 is shorter with the loss of *mer* operon.

The pangenome of the 1798 ST22 genomes consists of 4186 genes and the core genome consists of 2200 genes. Sequencing of ST22 is under-represented at a global level and previously the majority of the genomes have been sourced from samples originating in the European region. In this study adding 175 ST22 samples from India, we have filled-in some of the gaps in the global population. Analysis of the accessory genes with Panaroo has shown an increase in the number of accessory genes (cloud genes) from 1292 to 1406. This demonstrates the need for less biassed sequencing of samples collected from diverse geographical locations.

## Methods

### Bacterial isolates

A total of 509 random retrospective methicillin-resistant clinical *Staphylococcus aureus* isolates were obtained from 17 sentinel sites across 10 states of the country from the year 2014 to 2019 (Figure 1 & Table1) with the network established as previously described (44). Epidemiological metadata such as age, gender of patients, hospital, and infection type was collected from the sentinel sites. The laboratory assays were performed at the central hub for identification of S. aureus morphologically and performed Gram-Positive ID card on Vitek-2 compact system (Biomerieux). Based on the ID results, AST was tested using a P628 card, and MIC values were interpreted by WHONET & CLSI guidelines (45) (46) to characterize the isolate as Resistant (R) or Susceptible (S). Further, discordances between phenotypic antibiotic susceptibility profiles and WGS results from this study were analyzed to know the sensitivity and specificity of the test.

**Table 1:** A breakdown of the *S. aureus* Indian collection by clonal complexes.

| CC | Count |
| --- | --- |
| CC22 | 175 |
| CC8 | 103 |
| CC1 | 92 |
| Singletons | 56 |
| CC30 | 26 |
| CC5 | 23 |
| CC15 | 2 |
| CC45 | 1 |

**Table 2.**
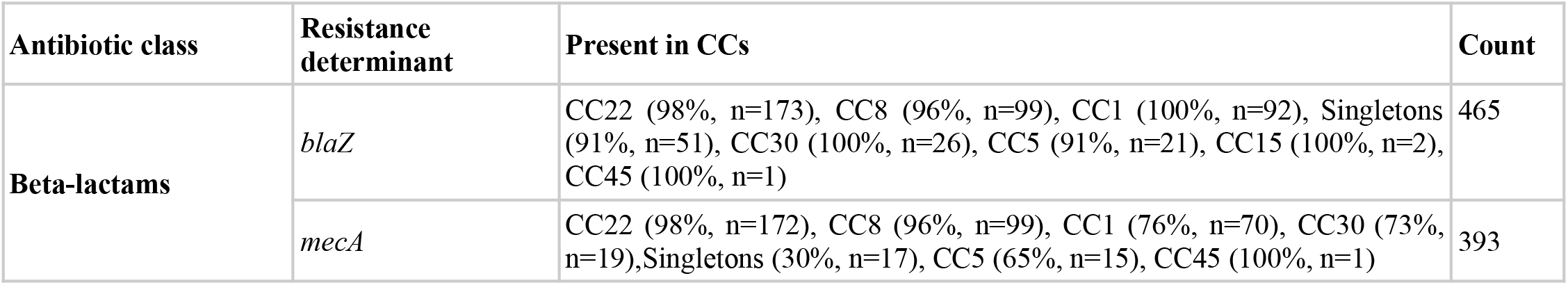
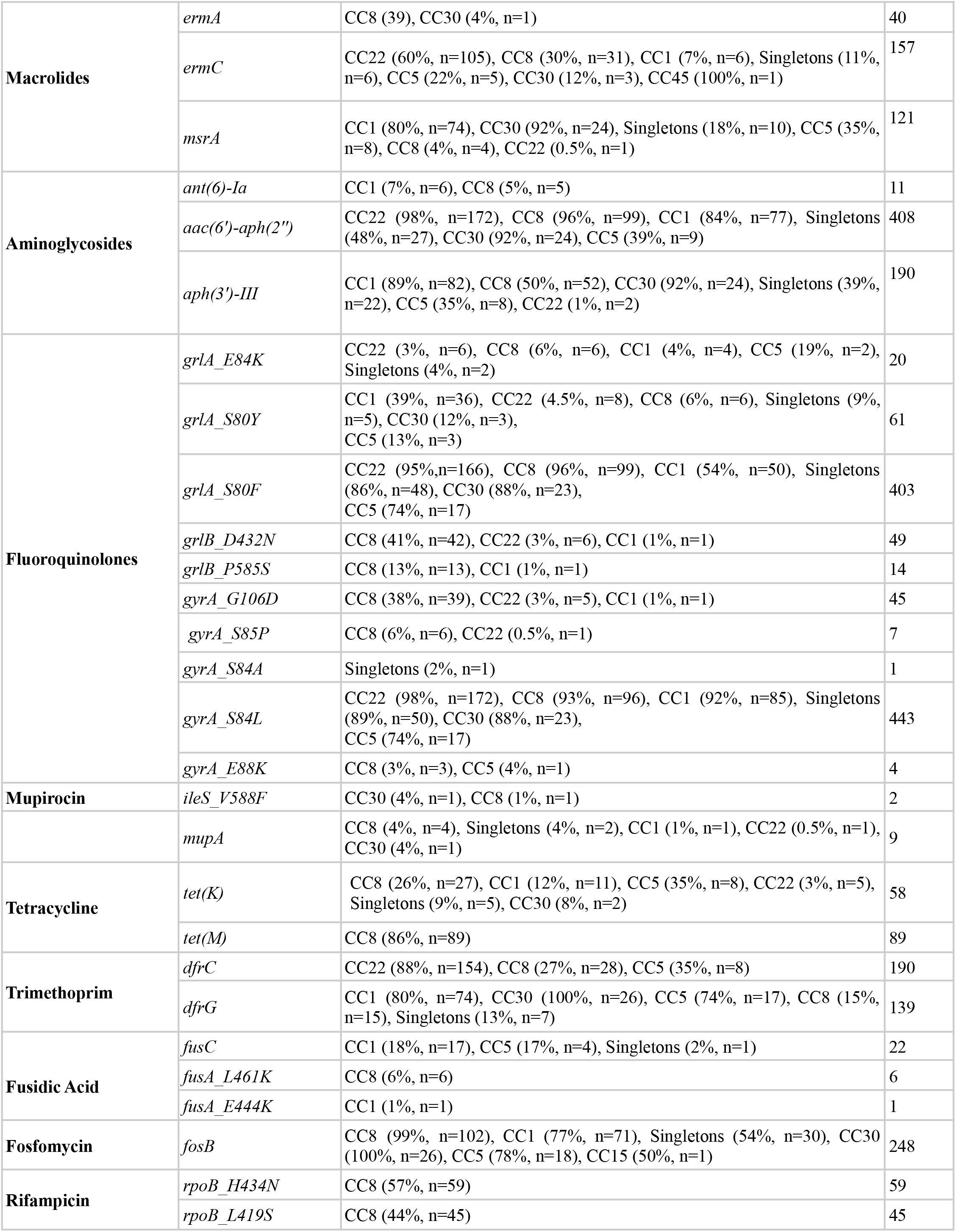
Number of isolates, per clonal complexes, carrying resistance determinants, associated with resistance to antibiotic classes.

### Whole-genome sequencing, assembly, and annotation

DNA was isolated using the QIAamp DNA mini kit and quantified using Qubit as instructed by the manufacturer. Genome libraries with 450 bp insert size were prepared and sequenced on the Illumina platform with paired-end reads of 150 bp length. The data were assembled using the Spades assembler v 3.14 (47) to generate contigs and annotated with Prokka v1.5 (48). Quality control of sequence data was performed using the GHRU QC pipeline based on (i) the basic statistics of raw reads, (ii) the assembly statistics, (iii) contamination due to SNV and sequences from different species, (iv) Species prediction using Bactinspector and (v) Overall QC as Pass, Warning or Fail of each isolate based on these different parameters.

### Variant detection and phylogenetic analysis

The 481 isolates were mapped to the reference genome *Staphylococcus aureus MSSA476* (strain GCF_000011525.1, ST1) using GHRU-SNP phylogeny pipeline v1.2.2 (49). The MGEs were masked in the pseudo genome alignment using MGEmasker (50) and the recombination region was removed using Gubbins algorithm v2.0.0 (51). The nonrecombinant SNPs were utilized to build a maximum-likelihood tree using IQ-tree (52) with parameters -czb to collapse near-zero branches, and a general time-reversible (GTR) model with 1000 bootstrap replicates. Visualization and phylogeographic analysis were performed on Microreact (53).

### In silico multilocus sequence typing

The PubMLST database was used to assign the sequence types to classify the diversity and epidemiology of isolates. The isolates were sequence typed with the MLST Oxford scheme using the 7 housekeeping genes (*arcC, aroE, glpF, gmk, pta, tpi, yqiL*) and assigned to an allele number. The allele numbers are combined to yield a specific sequence type (ST) utilizing the pubMLST database (https://pubmlst.org/organisms/staphylococcus-aureus). The goEburst algorithm (54) in PHYLOViZ (55) software (55) was applied to assign STs to Clonal Complexes (CCs). Those sharing identical alleles at six of seven loci were interpreted as single locus variants (SLVs).

### Novel MLST Characterisation

From the Ariba results the isolates with ST designated as novel or ST* were further analyzed using the R package MLSTar (56) as per the steps described in GHRU-Determining MLST allele sequences in novel STs (ref). The MLSTar queries the isolate sequence to reference the MLST database and gives a novel allele sequence or the profiles associated with each housekeeping gene. The novel profile or the novel allele sequences identified by MLSTar were submitted to PUBMLST and new STs were assigned.

### Antimicrobial resistance determinants and virulence genes

Genome data were analyzed for the presence of virulence genes using the virulence factor database (VFDB, (57)). Resfinder (58) was used to predict acquired AMR genes using the GHRU Resfinder pipeline (59)and mutations using the GHRU AMR pipeline (60) (with the PointFinder database (61). All the Bioinformatic analyses were performed using custom in-house developed nextflow pipelines as detailed on protocols.io (62).

### Plasmids

We ran Ariba v2.10.1 (63) with the PlasmidFinder database (64). We selected the 10 most prevalent plasmid replicon clusters in the datasets (GHRU isolates and CC22 global collection) and these are shown in the Microreact.

### Mobile Genetic Elements

SCC*mec* typing was performed on the assembled contigs using Staphopia (65). Staphopia includes a primer-based SCC*mec* Typing scheme, where the primers are aligned using BLAST. In many cases, Staphopia failed to distinguish between SCC*mec* type V and SCC*mec* type VII, because the identification of the SCC*mec* type is based on the *ccrC* genes. When there is more than one ccrC gene found for example in our samples the isolates had both *ccrC1* and *ccrC2* genes the Staphopia reports the sample as “SCC*mec* V VII”.This issue was resolved by uploading the genomes to SCC*mec*Finder (66). Some of the isolates were reported with multiple SCC*mec* types from Staphopia. These isolates were seen to carry extra SCC*mec* elements from other SCC*mec* cassettes. The spaTyper (67) was used to generate *spa* type identification for the isolates. The tool identifies the repeats and the order and generates a *spa* type. The repeat sequences and repeat orders found on http://spaserver2.ridom.de/ are used to identify the *spa* type of each enriched sequence. Ridom *spa* type and the genomic repeat sequence are then reported back to the user.

### Nanopore Sequencing and assembly

For the isolates for which the SCC*mec*finder gave an alert as “Additional complex(es) was found” and the ST772 isolates which were found to carry *mecA* but no SCC*mec* cassette was identified, we sequenced the representative isolates of these on Nanopore Minion-MK1C platform. The same extracted DNA used for the Illumina sequence was used with the rapid barcoding kit SQK-RBK004 on FLO-MIN106 R9 flow cells to perform long-read sequencing. High-accuracy base calling was done with guppy base caller(v4.3.4) used on minknow (v21.02.2) available at https://community.nanoporetech.com/downloads to get fastq files and the reads below q7 were filtered. Hybrid Assembly was performed with both Illumina and Nanopore reads using the nf-core Bacas assembly pipeline (68) (v1.1.1)(--assembler ‘unicycler’) to generate a single contig of the complete genome.

### Pan Genome analysis

The assembled contigs of 1798 ST22 isolates were annotated using prokka v1.5 (48). The gff files were given as the input to Panaroo (v1.2.8) (37) was used with “--clean-mode strict --remove-invalid-genes” to get the pangenome results.

### ST239 outbreak analysis

33 ST239 isolates collected from Bangalore between 2014 and 2017 and which formed a tight cluster in the *S. aureus* tree were inspected as a putative outbreak. A maximum-likelihood phylogenetic tree was generated for this cluster using *S. aureus* ST239 T0131 GCF_000204665.1_ASM20466v1 as a reference genome, identified based on similarity using BactInspector (69). Epidemiologic data, AST results, and genotypic characteristics of isolates of this cluster were then investigated.

### ST22 Global Isolates

For a number of 2 isolates, we could not get the exact isolation year. These were not included in the temporal analysis and are being shown with the year 2010, for visualization purposes only. (ERR234855, ERR234880). Temporal analysis was carried out using bactdating (35).

## Author contributions

This study was conceptualized by M.I.A., V.S., A.P. M.I.A performed the genomic analysis of a global collection of ST22 isolates. V.S. did the genomic analysis of the samples collected in this study. A.P. was involved in data summarization, table generation, and drafting of the manuscript. S.D.G. helped to resolve the SCC*mec* structures from the nanopore data. Funding for the study was provided through grants to K.L.R & D.A., V.G., G.N. & S.D. were involved in sample collection. V.G., S.D. & A.V. performed the microbiology part of the analysis & G.N. did the sequencing. A.U., S.D.G., S.A., reviewed the manuscript and suggested improvements.

## Conflicts of interest

The authors report no conflicts of interest. G.N., D.S., V.S., A.P., A.V., and K.L.R. report travel support from NIHR.

## Funding Statement

This work was supported by Official Development Assistance (ODA) funding from the National Institute for Health Research [grant number 16_136_111] and the Wellcome Trust grant number 206194.

This research was commissioned by the National Institute for Health Research using Official Development Assistance (ODA) funding. The views expressed in this publication are those of the authors and not necessarily those of the NHS, the National Institute for Health Research, or the Department of Health.

## Supplementary Figures

**Supplementary Figure 1:**
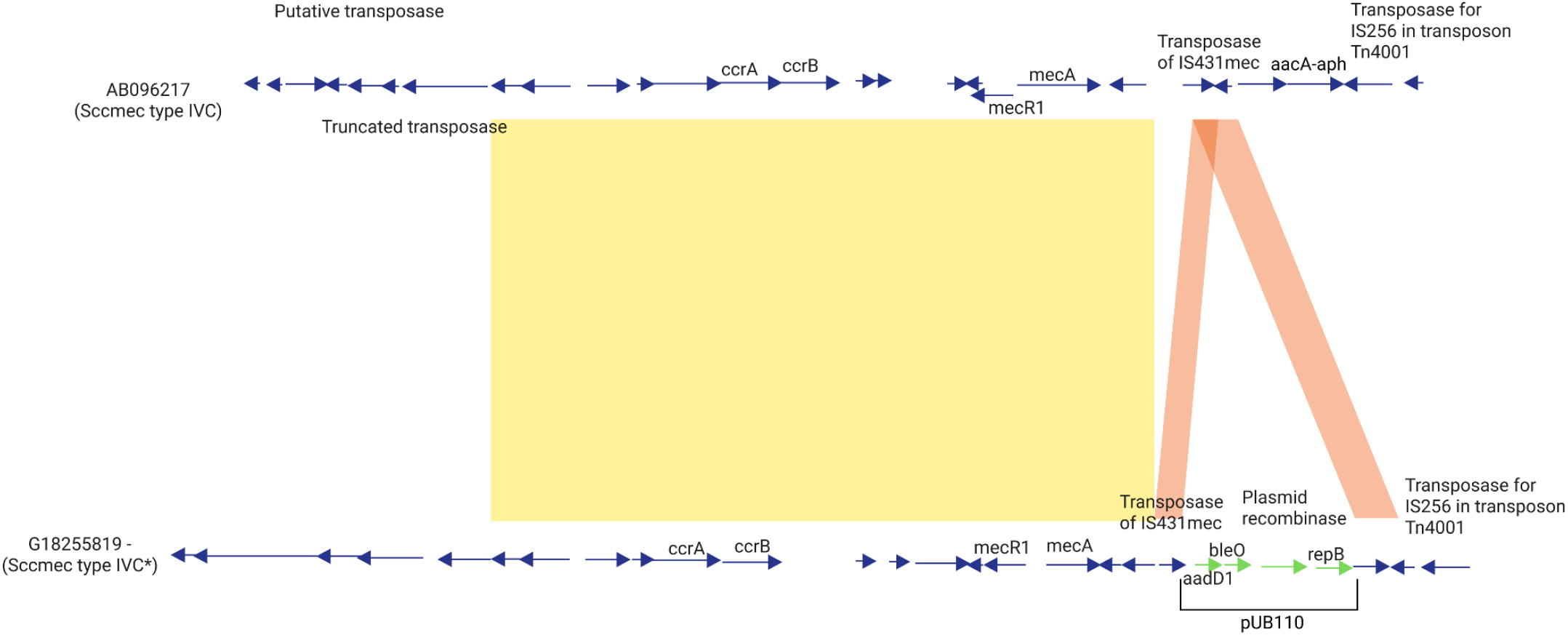
An ACT pairwise comparison between the SCC*mec* cassette found in isolate G18255819 (ST22, IND-1 Custer, marked as SCC*mec* type IVc in Microreact) and the template sequence AB096217.1 (SCC*mec* type IVc) shows that the IND-1 isolate has been integrated with pUB110 plasmid which carries the genes *aadD1* (conferring resistance to Aminoglycosides) and *bleO* (conferring resistance to Bleomycin) highlighted in green color.

**Supplementary Figure 2:**
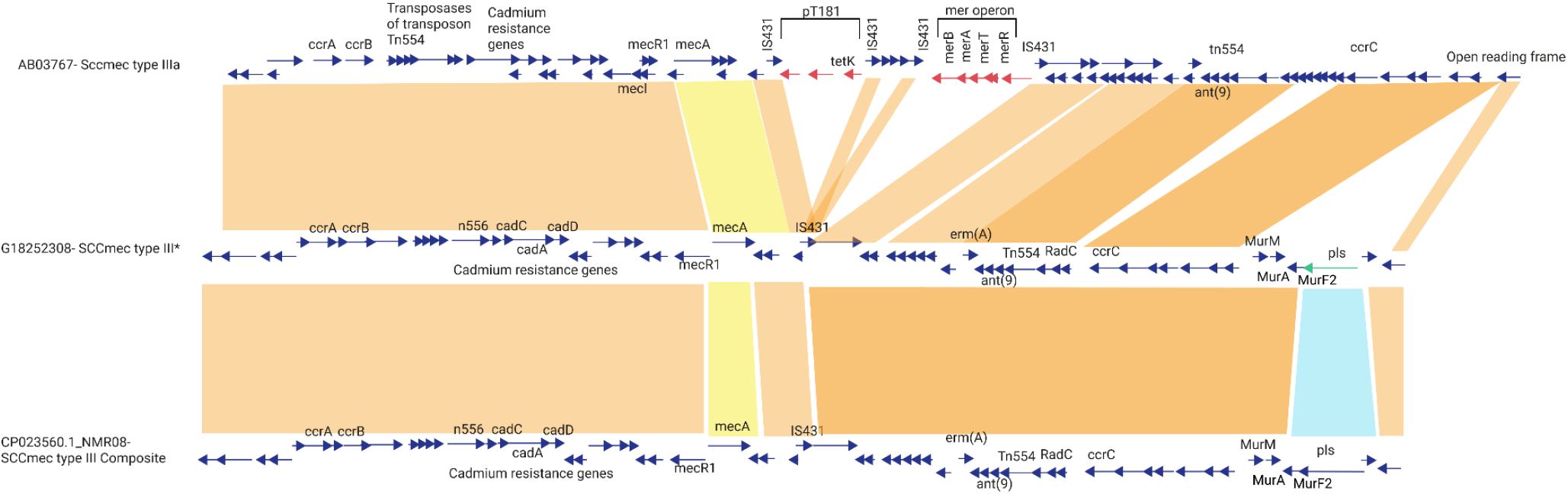
An ACT pairwise comparison between the SCC*mec* cassette found in an Indian isolate (G18252308, (ST239, SCC*mec* marked as “IIIa*” in the Microreact) and the template sequences AB037671 (SCC*mec* type IIIa) and CP023560.1_NMR08 (SCC*mec* type III composite) shows that the SCC*mec* type found in G18252308 has lost the mer operon & pT181. It has a variant of the *pls* gene which is highlighted in green color.

**Supplementary Figure 3:**
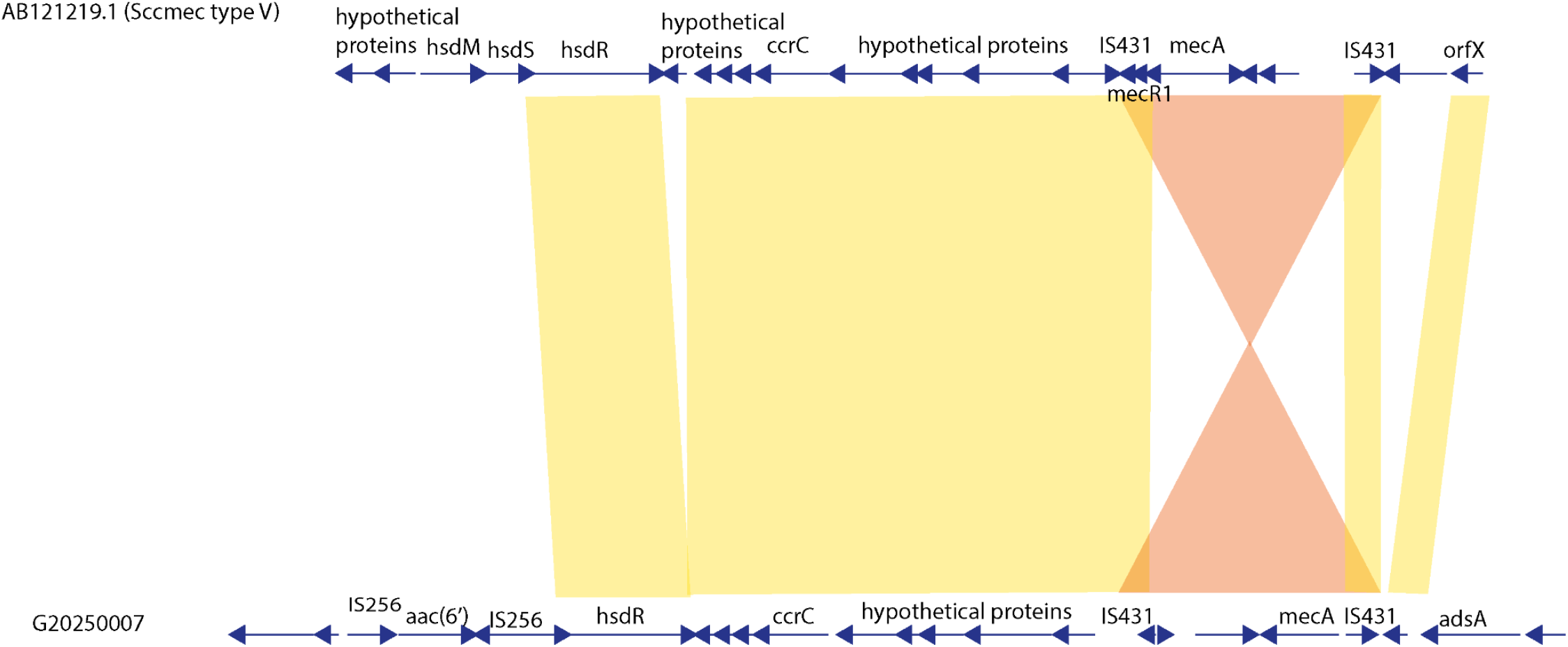
An ACT pairwise comparison between the SCC*mec* cassette found in isolate G20250007 (ST772, SCC*mec* cassette marked as type V* in Microreact) and the template sequence AB121219.1 (SCC*mec* type V) shows an insert sequence in the G20250007 cassette, carrying the *aac*(*6’*) gene, which confers resistance to Aminoglycosides.

## Notes

https://microreact.org/project/2xDvKQhriNveJ4kiVYsmSQ-s-aureus-wgs-study#i6az-ghru-tree

### Competing Interest Statement

The authors have declared no competing interest.

## References

1. WHO publishes list of bacteria for which new antibiotics are urgently needed [Internet]. [cited 2022 Jun 4]. Available from: https://www.who.int/news/item/27-02-2017-who-publishes-list-of-bacteria-for-which-new-antibiotics-are-urgently-needed

2. Loewen K, Schreiber Y, Kirlew M, Bocking N, Kelly L. Community-associated methicillin-resistant Staphylococcus aureus infection: Literature review and clinical update. Can Fam Physician [Internet]. 2017 Jul;63(7):512–20. Available from: https://www.ncbi.nlm.nih.gov/pubmed/28701438

3. Krishnan PU, Miles K, Shetty N. Detection of methicillin and mupirocin resistance in Staphylococcus aureus isolates using conventional and molecular methods: a descriptive study from a burns unit with high prevalence of MRSA. J Clin Pathol [Internet]. 2002 Oct;55(10):745–8. Available from: http://dx.doi.org/10.1136/jcp.55.10.745

4. Arakere G, Nadig S, Swedberg G, Macaden R, Amarnath SK, Raghunath D. Genotyping of methicillin-resistant Staphylococcus aureus strains from two hospitals in Bangalore, South India. J Clin Microbiol [Internet]. 2005 Jul;43(7):3198–202. Available from: http://dx.doi.org/10.1128/JCM.43.7.3198-3202.2005

5. Shambat S, Nadig S, Prabhakara S, Bes M, Etienne J, Arakere G. Clonal complexes and virulence factors of Staphylococcus aureus from several cities in India. BMC Microbiol [Internet]. 2012 May 1;12:64. Available from: http://dx.doi.org/10.1186/1471-2180-12-64

6. Goering RV, Shawar RM, Scangarella NE, O'Hara FP, Amrine-Madsen H, West JM, et al. Molecular epidemiology of methicillin-resistant and methicillin-susceptible Staphylococcus aureus isolates from global clinical trials. J Clin Microbiol [Internet]. 2008 Sep;46(9):2842–7. Available from: http://dx.doi.org/10.1128/JCM.00521-08

7. D'Souza N, Rodrigues C, Mehta A. Molecular characterization of methicillin-resistant Staphylococcus aureus with emergence of epidemic clones of sequence type (ST) 22 and ST 772 in Mumbai, India. J Clin Microbiol [Internet]. 2010 May;48(5):1806–11. Available from: http://dx.doi.org/10.1128/JCM.01867-09

8. Bouchiat C, El-Zeenni N, Chakrakodi B, Nagaraj S, Arakere G, Etienne J. Epidemiology of Staphylococcus aureus in Bangalore, India: emergence of the ST217 clone and high rate of resistance to erythromycin and ciprofloxacin in the community. New Microbes New Infect [Internet]. 2015 Sep;7:15–20. Available from: http://dx.doi.org/10.1016/j.nmni.2015.05.003

9. Sunagar R, Hegde NR, Archana GJ, Sinha AY, Nagamani K, Isloor S. Prevalence and genotype distribution of methicillin-resistant Staphylococcus aureus (MRSA) in India. J Glob Antimicrob Resist [Internet]. 2016 Dec;7:46–52. Available from: http://dx.doi.org/10.1016/j.jgar.2016.07.008

10. Dhawan B, Rao C, Udo EE, Gadepalli R, Vishnubhatla S, Kapil A. Dissemination of methicillin-resistant Staphylococcus aureus SCCmec type IV and SCCmec type V epidemic clones in a tertiary hospital: challenge to infection control. Epidemiol Infect [Internet]. 2015 Jan;143(2):343–53. Available from: http://dx.doi.org/10.1017/S095026881400065X

11. Bakthavatchalam YD, Vasudevan K, Rao S, Varughese S, Rupali P, Gina M, et al. Genomic portrait of community-associated methicillin-resistant Staphylococcus aureus ST772-SCCmec V lineage from India. Gene Reports [Internet]. 2021 Sep 1;24:101235. Available from: https://www.sciencedirect.com/science/article/pii/S245201442100220X

12. Rajan V, Schoenfelder SMK, Ziebuhr W, Gopal S. Genotyping of community-associated methicillin resistant Staphylococcus aureus (CA-MRSA) in a tertiary care centre in Mysore, South India: ST2371-SCCmec IV emerges as the major clone. Infect Genet Evol [Internet]. 2015 Aug;34:230–5. Available from: http://dx.doi.org/10.1016/j.meegid.2015.05.032

13. Boakes E, Kearns AM, Ganner M, Perry C, Hill RL, Ellington MJ. Distinct bacteriophages encoding Panton-Valentine leukocidin (PVL) among international methicillin-resistant Staphylococcus aureus clones harboring PVL. J Clin Microbiol [Internet]. 2011 Feb;49(2):684–92. Available from: http://dx.doi.org/10.1128/JCM.01917-10

14. Prabhakara S, Khedkar S, Loganathan RM, Chandana S, Gowda M, Arakere G, et al. Draft genome sequence of Staphylococcus aureus 118 (ST772), a major disease clone from India. J Bacteriol [Internet]. 2012 Jul;194(14):3727–8. Available from: http://dx.doi.org/10.1128/JB.00480-12

15. Steinig EJ, Duchene S, Robinson DA, Monecke S, Yokoyama M, Laabei M, et al. Evolution and Global Transmission of a Multidrug-Resistant, Community-Associated Methicillin-Resistant Staphylococcus aureus Lineage from the Indian Subcontinent. MBio [Internet]. 2019 Nov 26;10(6). Available from: http://dx.doi.org/10.1128/mBio.01105-19

16. Monecke S, Baier V, Coombs GW, Slickers P, Ziegler A, Ehricht R. Genome sequencing and molecular characterisation of Staphylococcus aureus ST772-MRSA-V, "Bengal Bay Clone." BMC Res Notes [Internet]. 2013 Dec 20;6:548. Available from: http://dx.doi.org/10.1186/1756-0500-6-548

17. Steinig EJ, Andersson P, Harris SR, Sarovich DS, Manoharan A, Coupland P, et al. Single-molecule sequencing reveals the molecular basis of multidrug-resistance in ST772 methicillin-resistant Staphylococcus aureus. BMC Genomics [Internet]. 2015 May 16;16:388. Available from: http://dx.doi.org/10.1186/s12864-015-1599-9

18. Enright MC, Robinson DA, Randle G, Feil EJ, Grundmann H, Spratt BG. The evolutionary history of methicillin-resistant Staphylococcus aureus (MRSA). Proc Natl Acad Sci U S A [Internet]. 2002 May 28;99(11):7687–92. Available from: http://dx.doi.org/10.1073/pnas.122108599

19. Feil EJ, Nickerson EK, Chantratita N, Wuthiekanun V, Srisomang P, Cousins R, et al. Rapid detection of the pandemic methicillin-resistant Staphylococcus aureus clone ST 239, a dominant strain in Asian hospitals. J Clin Microbiol [Internet]. 2008 Apr;46(4):1520–2. Available from: http://dx.doi.org/10.1128/JCM.02238-07

20. De Backer S, Xavier BB, Vanjari L, Coppens J, Lammens C, Vemu L, et al. Remarkable geographical variations between India and Europe in carriage of the staphylococcal surface protein-encoding sasX/sesI and in the population structure of methicillin-resistant Staphylococcus aureus belonging to clonal complex 8. Clin Microbiol Infect [Internet]. 2019 May;25(5):628.e1–628.e7. Available from: http://dx.doi.org/10.1016/j.cmi.2018.07.024

21. Neetu TJP, Murugan S. Genotyping of Methicillin Resistant Staphylococcus aureus from Tertiary Care Hospitals in Coimbatore, South India. J Glob Infect Dis [Internet]. 2016 Apr;8(2):68–74. Available from: http://dx.doi.org/10.4103/0974-777X.182119

22. Abimanyu N, Murugesan S, Krishnan P. Emergence of methicillin-resistant Staphylococcus aureus ST239 with high-level mupirocin and inducible clindamycin resistance in a tertiary care center in Chennai, South India. J Clin Microbiol [Internet]. 2012 Oct;50(10):3412–3. Available from: http://dx.doi.org/10.1128/JCM.01663-12

23. Holden MTG, Hsu LY, Kurt K, Weinert LA, Mather AE, Harris SR, et al. A genomic portrait of the emergence, evolution, and global spread of a methicillin-resistant Staphylococcus aureus pandemic. Genome Res [Internet]. 2013 Apr;23(4):653–64. Available from: http://dx.doi.org/10.1101/gr.147710.112

24. Biber A, Abuelaish I, Rahav G, Raz M, Cohen L, Valinsky L, et al. A typical hospital-acquired methicillin-resistant Staphylococcus aureus clone is widespread in the community in the Gaza strip. PLoS One [Internet]. 2012 Aug 16;7(8):e42864. Available from: http://dx.doi.org/10.1371/journal.pone.0042864

25. Gostev V, Leyn S, Kruglov A, Likholetova D, Kalinogorskaya O, Baykina M, et al. Global Expansion of Linezolid-Resistant Coagulase-Negative Staphylococci. Front Microbiol [Internet]. 2021;12. Available from: https://www.frontiersin.org/article/10.3389/fmicb.2021.661798

26. Nadig S, Ramachandra Raju S, Arakere G. Epidemic meticillin-resistant Staphylococcus aureus (EMRSA-15) variants detected in healthy and diseased individuals in India. J Med Microbiol [Internet]. 2010 Jul;59(Pt 7):815–21. Available from: http://dx.doi.org/10.1099/jmm.0.017632-0

27. Alfouzan W, Udo EE, Modhaffer A, Alosaimi A 'a. Molecular Characterization of Methicillin-Resistant Staphylococcus aureus in a Tertiary Care hospital in Kuwait. Sci Rep [Internet]. 2019 Dec 6;9(1):18527. Available from: http://dx.doi.org/10.1038/s41598-019-54794-8

28. Pokhrel RH, Aung MS, Thapa B, Chaudhary R, Mishra SK, Kawaguchiya M, et al. Detection of ST772 Panton-Valentine leukocidin-positive methicillin-resistant Staphylococcus aureus (Bengal Bay clone) and ST22 S. aureus isolates with a genetic variant of elastin binding protein in Nepal. New Microbes New Infect [Internet]. 2016 May;11:20–7. Available from: http://dx.doi.org/10.1016/j.nmni.2016.02.001

29. Yamaguchi T, Nakamura I, Chiba K, Matsumoto T. Epidemiological and microbiological analysis of community-associated methicillin-resistant Staphylococcus aureus strains isolated from a Japanese hospital. Jpn J Infect Dis [Internet]. 2012;65(2):175–8. Available from: https://www.ncbi.nlm.nih.gov/pubmed/22446128

30. Xiao N, Yang J, Duan N, Lu B, Wang L. Community-associated Staphylococcus aureus PVL+ ST22 predominates in skin and soft tissue infections in Beijing, China. Infect Drug Resist [Internet]. 2019 Aug 12;12:2495–503. Available from: http://dx.doi.org/10.2147/IDR.S212358

31. Senok A, Somily A, Raji A, Gawlik D, Al-Shahrani F, Baqi S, et al. Diversity of methicillin-resistant Staphylococcus aureus CC22-MRSA-IV from Saudi Arabia and the Gulf region. Int J Infect Dis [Internet]. 2016 Oct;51:31–5. Available from: http://dx.doi.org/10.1016/j.ijid.2016.08.016

32. Coppens J, Xavier BB, Vanjari L, De Backer S, Lammens C, Vemu L, et al. Novel composite SCCmec type III element in ST239 MRSA isolated from an Indian hospital. J Antimicrob Chemother [Internet]. 2019 Jan 1;74(1):264–6. Available from: http://dx.doi.org/10.1093/jac/dky399

33. Aravind P, Krishnan PU, Srinivasa H, Joseph V. Screening of Burns Unit staff of a tertiary care hospital for methicillin-resistant staphylococcus aureus colonisation. Mcgill J Med [Internet]. 2020 Dec 1;5(2). Available from: https://mjm.mcgill.ca/article/view/701

34. Harris SR, Cartwright EJP, Török ME, Holden MTG, Brown NM, Ogilvy-Stuart AL, et al. Whole-genome sequencing for analysis of an outbreak of meticillin-resistant Staphylococcus aureus: a descriptive study. Lancet Infect Dis [Internet]. 2013 Feb;13(2):130–6. Available from: http://dx.doi.org/10.1016/S1473-3099(12)70268-2

35. Didelot X, Croucher NJ, Bentley SD, Harris SR, Wilson DJ. Bayesian inference of ancestral dates on bacterial phylogenetic trees. Nucleic Acids Res [Internet]. 2018 Dec 14;46(22):e134. Available from: http://dx.doi.org/10.1093/nar/gky783

36. Gostev V, Ivanova K, Kruglov A, Kalinogorskaya O, Ryabchenko I, Zyryanov S, et al. Comparative genome analysis of global and Russian strains of community-acquired methicillin-resistant Staphylococcus aureus ST22, a "Gaza clone." Int J Antimicrob Agents [Internet]. 2021 Feb;57(2):106264. Available from: http://dx.doi.org/10.1016/j.ijantimicag.2020.106264

37. Tonkin-Hill G, MacAlasdair N, Ruis C, Weimann A, Horesh G, Lees JA, et al. Producing polished prokaryotic pangenomes with the Panaroo pipeline. Genome Biol [Internet]. 2020 Jul 22;21(1):180. Available from: http://dx.doi.org/10.1186/s13059-020-02090-4

38. Siboo IR, Chambers HF, Sullam PM. Role of SraP, a Serine-Rich Surface Protein of Staphylococcus aureus, in binding to human platelets. Infect Immun [Internet]. 2005 Apr;73(4):2273–80. Available from: http://dx.doi.org/10.1128/IAI.73.4.2273-2280.2005

39. Di Gregorio S, Haim MS, Vielma Vallenilla J, Cohen V, Rago L, Gulone L, et al. Genomic Epidemiology of CC30 Methicillin-Resistant Staphylococcus aureus Strains from Argentina Reveals Four Major Clades with Distinctive Genetic Features. mSphere [Internet]. 2021 Mar 10;6(2). Available from: http://dx.doi.org/10.1128/mSphere.01297-20

40. Toleman MS, Reuter S, Coll F, Harrison EM, Blane B, Brown NM, et al. Systematic Surveillance Detects Multiple Silent Introductions and Household Transmission of Methicillin-Resistant Staphylococcus aureus USA300 in the East of England. J Infect Dis [Internet]. 2016 Aug 1;214(3):447–53. Available from: http://dx.doi.org/10.1093/infdis/jiw166

41. Tamarapu S, McKillip JL, Drake M. Development of a multiplex polymerase chain reaction assay for detection and differentiation of Staphylococcus aureus in dairy products. J Food Prot [Internet]. 2001 May;64(5):664–8. Available from: http://dx.doi.org/10.4315/0362-028x-64.5.664

42. Novick RP. Mobile genetic elements and bacterial toxinoses: the superantigen-encoding pathogenicity islands of Staphylococcus aureus. Plasmid [Internet]. 2003 Mar;49(2):93–105. Available from: http://dx.doi.org/10.1016/s0147-619x(02)00157-9

43. Health Organization. Implementation of Early Warning and Response with a focus on Event-Based Surveillance. Early Detect Assess response to acute public Heal.

44. Nagaraj G, Shamanna V, Govindan V, Rose S, Sravani D, Akshata KP, et al. High-Resolution Genomic Profiling of Carbapenem-Resistant Klebsiella pneumoniae Isolates: A Multicentric Retrospective Indian Study. Clin Infect Dis [Internet]. 2021 Dec 1;73(Suppl_4):S300–7. Available from: http://dx.doi.org/10.1093/cid/ciab767

45. Stelling JM, O'Brien TF. Surveillance of antimicrobial resistance: the WHONET program. Clin Infect Dis [Internet]. 1997 Jan;24 Suppl 1:S157–68. Available from: http://dx.doi.org/10.1093/clinids/24.supplement_1.s157

46. Weinstein, Limbago, Patel, Mathers. M100 performance standards for antimicrobial susceptibility testing. & Laboratory Standards….

47. Prjibelski A, Antipov D, Meleshko D, Lapidus A, Korobeynikov A. Using SPAdes De Novo Assembler. Curr Protoc Bioinformatics [Internet]. 2020 Jun;70(1):e102. Available from: http://dx.doi.org/10.1002/cpbi.102

48. Seemann T. Prokka: rapid prokaryotic genome annotation. Bioinformatics [Internet]. 2014 Jul 15;30(14):2068–9. Available from: http://dx.doi.org/10.1093/bioinformatics/btu153

49. Snp_phylogeny [Internet]. GitLab. [cited 2022 Jun 4]. Available from: https://gitlab.com/cgps/ghru/pipelines/snp_phylogeny

50. MGEmasker [Internet]. GitLab. [cited 2022 Jun 4]. Available from: https://gitlab.com/antunderwood/mgemasker

51. Croucher NJ, Page AJ, Connor TR, Delaney AJ, Keane JA, Bentley SD, et al. Rapid phylogenetic analysis of large samples of recombinant bacterial whole genome sequences using Gubbins. Nucleic Acids Res [Internet]. 2015 Feb 18;43(3):e15. Available from: http://dx.doi.org/10.1093/nar/gku1196

52. Nguyen LT, Schmidt HA, von Haeseler A, Minh BQ. IQ-TREE: a fast and effective stochastic algorithm for estimating maximum-likelihood phylogenies. Mol Biol Evol [Internet]. 2015 Jan;32(1):268–74. Available from: http://dx.doi.org/10.1093/molbev/msu300

53. Argimón S, Abudahab K, Goater RJE, Fedosejev A, Bhai J, Glasner C, et al. Microreact: visualizing and sharing data for genomic epidemiology and phylogeography. Microb Genom [Internet]. 2016 Nov;2(11):e000093. Available from: http://dx.doi.org/10.1099/mgen.0.000093

54. Francisco AP, Bugalho M, Ramirez M, Carriço JA. Global optimal eBURST analysis of multilocus typing data using a graphic matroid approach. BMC Bioinformatics [Internet]. 2009 May 18;10:152. Available from: http://dx.doi.org/10.1186/1471-2105-10-152

55. Nascimento M, Sousa A, Ramirez M, Francisco AP, Carriço JA, Vaz C. PHYLOViZ 2.0: providing scalable data integration and visualization for multiple phylogenetic inference methods. Bioinformatics [Internet]. 2017 Jan 1;33(1):128–9. Available from: http://dx.doi.org/10.1093/bioinformatics/btw582

56. Ferrés I, Iraola G. MLSTar: automatic multilocus sequence typing of bacterial genomes in R. PeerJ [Internet]. 2018 Jun 15;6:e5098. Available from: http://dx.doi.org/10.7717/peerj.5098

57. Liu B, Zheng D, Jin Q, Chen L, Yang J. VFDB 2019: a comparative pathogenomic platform with an interactive web interface. Nucleic Acids Res [Internet]. 2019 Jan 8;47(D1):D687–92. Available from: http://dx.doi.org/10.1093/nar/gky1080

58. Bortolaia V, Kaas RS, Ruppe E, Roberts MC, Schwarz S, Cattoir V, et al. ResFinder 4.0 for predictions of phenotypes from genotypes. J Antimicrob Chemother [Internet]. 2020 Dec 1;75(12):3491–500. Available from: http://dx.doi.org/10.1093/jac/dkaa345

59. Resfinder-4 [Internet]. GitLab. [cited 2022 Jun 4]. Available from: https://gitlab.com/cgps/ghru/pipelines/dsl2/pipelines/resfinder-4

60. Amr_prediction [Internet]. GitLab. [cited 2022 Jun 4]. Available from: https://gitlab.com/cgps/ghru/pipelines/dsl2/pipelines/amr_prediction

61. Zankari E, Allesøe R, Joensen KG, Cavaco LM, Lund O, Aarestrup FM. PointFinder: a novel web tool for WGS-based detection of antimicrobial resistance associated with chromosomal point mutations in bacterial pathogens. J Antimicrob Chemother [Internet]. 2017 Oct 1;72(10):2764–8. Available from: http://dx.doi.org/10.1093/jac/dkx217

62. Underwood A. GHRU (Genomic Surveillance of antimicrobial resistance) retrospective 1 bioinformatics methods [Internet]. protocols.io. 2020 [cited 2022 Jun 4]. Available from: https://www.protocols.io/view/ghru-genomic-surveillance-of-antimicrobial-resista-bpn6mmhe

63. Hunt M, Mather AE, Sánchez-Busó L, Page AJ, Parkhill J, Keane JA, et al. ARIBA: rapid antimicrobial resistance genotyping directly from sequencing reads. Microb Genom [Internet]. 2017 Oct;3(10):e000131. Available from: http://dx.doi.org/10.1099/mgen.0.000131

64. Carattoli A, Hasman H. PlasmidFinder and In Silico pMLST: Identification and Typing of Plasmid Replicons in Whole-Genome Sequencing (WGS). Methods Mol Biol [Internet]. 2020;2075:285–94. Available from: http://dx.doi.org/10.1007/978-1-4939-9877-7_20

65. Petit RA 3rd, Read TD. Staphylococcus aureus viewed from the perspective of 40,000+ genomes. PeerJ [Internet]. 2018 Jul 12;6:e5261. Available from: http://dx.doi.org/10.7717/peerj.5261

66. Kaya H, Hasman H, Larsen J, Stegger M, Johannesen TB, Allesøe RL, et al. SCCmecFinder, a Web-Based Tool for Typing of Staphylococcal Cassette Chromosome mec in Staphylococcus aureus Using Whole-Genome Sequence Data. mSphere [Internet]. 2018 Jan;3(1). Available from: http://dx.doi.org/10.1128/mSphere.00612-17

67. Sanchez-Herrero JF, mjsull. spaTyper: Staphylococcal protein A (spa) characterization pipeline [Internet]. 2020. Available from: https://zenodo.org/record/4063625

68. Peltzer A, Straub D, Bot NC, Garcia MU, Taylor B, Angelov A, et al. nf-core/bacass: v2.0.0 nf-core/bacass: "Navy Steel Swordfish" [Internet]. 2021. Available from: https://zenodo.org/record/5289278

69. BactInspector [Internet]. GitLab. [cited 2022 Jun 4]. Available from: https://gitlab.com/antunderwood/bactinspector

